# Resting-state functional connectivity and psychopathology in Klinefelter syndrome (47, XXY)

**DOI:** 10.1101/2020.12.15.422868

**Authors:** Ethan T. Whitman, Siyuan Liu, Erin Torres, Allysa Warling, Kathleen Wilson, Ajay Nadig, Cassidy McDermott, Liv S. Clasen, Jonathan D. Blumenthal, François M. Lalonde, Stephen J. Gotts, Alex Martin, Armin Raznahan

## Abstract

Klinefelter syndrome (47, XXY; Henceforth: XXY syndrome) is a high impact but poorly understood genetic risk factor for neuropsychiatric impairment. Here, we provide the first neuroimaging study to map resting-state functional connectivity (rsFC) changes in XXY syndrome and ask how these might relate to brain anatomy and psychopathology. We collected resting state functional magnetic resonance imaging data from 75 individuals with XXY and 84 healthy XY males. We implemented a brain-wide screen to identify regions with altered global rsFC in XXY vs. XY males, and then used seed-based analysis to decompose these alterations. We further compared rsFC changes with regional changes in brain volume from voxel-based morphometry and tested for correlations between rsFC and symptom variation within XXY syndrome. We found that XXY syndrome was characterized by increased global rsFC in the left dorsolateral prefrontal cortex (DLPFC), associated with overconnectivity with diverse rsFC networks. Regional rsFC changes were partly coupled to regional volumetric changes in XXY syndrome. Within the precuneus, variation in DLPFC rsFC within XXY syndrome was correlated with the severity of psychopathology in XXY individuals. Our findings provide the first view of altered functional brain connectivity in XXY syndrome and delineate links between these alterations and those relating to both brain anatomy and psychopathology. Taken together, these insights advance biological understanding of XXY syndrome as a disorder in its own right, and as a model of genetic risk for psychopathology more broadly.

## INTRODUCTION

Klinefelter syndrome (47, XXY karyotype) is a sex chromosome aneuploidy disorder estimated to occur in approximately 1 in 633 male births [1]. This aneuploidy is associated with increased rates of neuropsychiatric disorders including autism, attention-deficit/hyperactivity disorder, and mood disorders relative to karyotypically normal males [2–5]. XXY syndrome increases risk for language difficulties alongside a modest decrease in general cognitive ability [2]. The biological bases of neuropsychiatric risk in XXY syndrome remain poorly understood, but there is extensive evidence for reproducible changes in regional brain anatomy from in vivo structural neuroimaging data. These changes include decreased total brain volume with relative contractions in temporal, frontal, and cerebellar structures and relative expansion in parietal-occipital regions [6–8]. The presence of overlapping regional anatomy changes in other disorders of X-chromosome dosage (e.g. trisomy X [9, 10]; and Turner syndrome [9, 10]) suggests a direct role for X-linked genes in driving the observed anatomical brain changes in XXY syndrome.

Although there is a consensus regarding the spatial pattern of regional neuroanatomical changes in XXY syndrome, functional brain organization in XXY remains largely unstudied [11]. In contrast to several genetically defined neuropsychiatric disorders that are less common than XXY syndrome [e.g. 22q11.2 deletions [12–15], 16p11.2 copy number variations [13, 16], Fragile X disorder [17]], we still lack a brain-wide study of resting-state functional connectivity (rsFC) in XXY syndrome. Our study seeks to address this gap in knowledge with two main considerations in mind.

First, from the perspective of psychiatric genetics, XXY syndrome offers a powerful model for “genetics-first” research [18] into mechanisms of biological risk for psychopathology. Our study empirically considers three general questions regarding mechanisms of genetic risk in the human brain that remain largely unaddressed to date: (i) the degree to which genetically driven disruptions of brain structure and function are colocalized or differentially distributed across the brain [19], (ii) the degree to which the considerable variation in neuropsychiatric impairments within a neurogenetic disorder can be related to inter-individual variation in rsFC [20, 21], and (iii) whether any such rsFC-behavior relationships amongst patients are colocalized with regions of altered rFC between patients and controls.

Second, the fact that XXY syndrome models X-chromosome dosage effects in humans means that better understanding brain changes in XXY syndrome would help address broader questions relating to X-chromosome biology. Specifically, because X-chromosome dosage disparity is a foundational genetic difference between males and females, X-chromosome dosage effects are candidate contributors to sex differences in brain structure and function [22–27]. Furthermore, modelling X-chromosome dosage effects on rsFC through the study of XXY syndrome may help interpreting past observations that X-linked genes show elevated levels of brain expression relative to Y-linked and autosomal genes [28, 29] and an enrichment for genetic variants associated with intellectual disability syndromes [28, 29].

Motivated by these factors, we used resting-state functional magnetic resonance imaging (fMRI) to study rsFC in 75 males with XXY and 84 age-matched XY male controls. Our analytic approach first provides a brain-wide screen for regions with altered global rsFC in XXY, and then characterizes the patterns of interregional dysconnectivity that drive these effects [30]. We also test whether the spatial distribution of rsFC changes is related to that of regional brain volume changes. Finally, we examine whether the marked variability in neuropsychiatric outcome severity within XXY syndrome is related to variability in rsFC, and whether any such relationships align with rsFC changes in XXY syndrome as compared to typically developing XY controls.

## MATERIALS AND METHODS

### Participants

Participants were recruited to participate in this study through XXY parent support organizations, the National Institute of Mental Health (NIMH) website, and the National Institutes of Health (NIH) healthy volunteer office. The sample comprised 75 XXY participants (ages 6-25, mean age 16.7 years) and 84 XY controls (ages 7-25, mean age 17.2 years). All participants had normal radiological reports and no participants had a history of brain injury or comorbid neurological disorders. All XXY participants were non-mosaic with a diagnosis by karyotype, and all XY participants were screened to exclude a history of neurodevelopmental or psychiatric disorders. This study was approved by the NIH Combined Neuroscience Institutional Review Board. All participants gave consent or assent, as appropriate. All protocols were completed at the NIH Clinical Center in Bethesda, Maryland.

### Cognitive and Behavioral measures

Full-scale Intelligence Quotient (henceforth “IQ”) was measured for the XXY group using the Wechsler Intelligence Scale for Children, Fifth Edition (WISC-V) and the Wechsler Adult Intelligence Scale, Fourth Edition (WAIS-IV; [31]). Global total psychopathology was measured using age-normed t-scores from the Child Behavior Checklist (CBCL; [32]) and the Adult Behavior Checklist (ABCL; [33]).

### Neuroimaging

Neuroimaging data were collected using a MR750 3-Tesla (General Electric, Wisconsin) whole-body scanner with a 32-channel head coil. High-resolution MP-RAGE T1-weighted anatomical images were collected for each participant (176 contiguous sagittal slices with 256 x 256 in-plane matrix and 1mm slice thickness yielding 1mm isotropic voxels). Spontaneous slowly fluctuating brain activity was measured during fMRI through an echo-planar imaging (EPI) sequence while participants were instructed to lay still and focus on a fixation cross for 10 minutes (repetition time of 2 seconds, echo time = 30ms, flip angle = 60 degrees, 41 interleaved axial slices per volume, 3mm slice thickness in a 216mm x 216mm acquisition matrix, single voxel = 3mm isotropic). Resting-state data were evaluated for transient head motion artifacts using the AFNI program @1dDiffMag [34] to quantify the magnitude of head motion throughout the scan (mm/repetition time [TR]) comparable to average framewise displacement [35, 36]. Only participants with complete resting-state scans and average head motion < 0.3mm/TR were included.

### Preprocessing and analysis of resting-state fMRI data

Preprocessing was performed using the AFNI software package [34] based on the ANATICOR approach [37]. We removed physiological noise from the echo-planar data by performing principal component analysis (PCA) to identify the first three principal components within ventricles and white matter. We used linear regression to remove these three primary components, head motion, average signal from ventricles, and a local average of white matter signal within a 20mm radius centered on each voxel. We then smoothed EPI data with a 6.0mm Gaussian kernel and non-linearly transformed scans into standard Talairach space [40] (for more detail see **Supplementary Text 1**).

We calculated voxel-wise global rsFC maps for each participant by finding the average correlation of the EPI data at one voxel with the time series at every other voxel in the gray matter mask (as described in [30]). For voxels present in 90% of individual-participant grey matter masks, we Fisher’s z-transformed the voxel-wise correlation coefficients and compared groups using two-sample t-tests at each voxel while covarying for age and average head motion. We corrected for multiple comparisons across voxels by applying a voxel-level p < 0.002 threshold, and a cluster size threshold of 21 -- thereby achieving a corrected FWE of 5% within the revised version of AFNI’s 3dClustSim function, after the bug fix in 2017 [41]. This identified two significant clusters of increased global connectivity in XXY syndrome: a 49-voxel cluster in the left DLPFC and a 22-voxel cluster in the right medial temporal lobe.

To ensure the stability of any group difference clusters detected and to eliminate any possibility of noise bias for subsequent seed tests, we implemented a leave-one-out validation technique by recalculating group differences 159 times while iteratively excluding one participant at a time [42]. We identified any clusters that were statistically significant in all 159 iterations as stable clusters of group difference in global connectivity. This procedure --which screened all gray matter voxels to identify foci with statistically significant and reproducible differences in average rsFC with other brain regions in XXY vs. XY groups -identified a 32-voxel region within the aforementioned left DLPFC cluster showing consistent and significantly increased global connectivity in XXY. By definition, this region was detected in a robust manner that did not depend on noise present in any of the individual datasets. The cluster detected in the medial temporal lobe did not survive leave-one-out validation and was excluded from subsequent analyses.

Next, a seed-based analysis was used to identify the specific patterns of altered inter-regional connectivity that contributed to the observed 32-voxel cluster of increased DLPFC global connectivity in XXY syndrome. These seed analyses were considered to be statistically independent from the prior screen for altered global rsFC because of the leave-one-out criterion that has been used to define the seed [43]). For each individual, we correlated average EPI time series within the left DLPFC and every other gray matter voxel. Then, after Fisher’s z-transformation, we performed two-sample t-tests between groups at each voxel, co-varying for participant age and average head motion. This identified the set of target regions which showed altered rsFC with the robust left DLPFC seed in XXY vs. XY groups after correction using the revised version of AFNI’s 3dClustsim to ensure FWE of 5% [41, see above]. Target regions were parcellated using the boundaries along major anatomical divisions and remaining large clusters exceeding the 85th centile of target cluster size distribution were further divided using a previously-published rsFC parcellation of the brain [44]. This process yielded 39 regions of interest (ROIs), including the original DLPFC seed region. Group differences in rsFC between all unique ROI pairs (i.e. rsFC “edges”) were determined using two-sample t-tests between groups. We considered edges to be statistically significant if they had absolute t-scores above the 95th percentile of an empirical null distribution formed from the most extreme edge-level t-scores from 1000 permutations of group assignment.

### Comparing rsFC and anatomical alterations in XXY

Voxel-wise group differences in gray matter volume (GMV) between XXY and XY groups were estimated using the DARTEL procedure in SPM 12. Images were corrected for magnetic field inhomogeneity, segmented, and normalized to the Montreal Neurological Institute (MNI) space in a unified model. Gray matter probability maps were generated for each individual and smoothed. To find group differences in regional GMV, a voxel-wise ANOVA model in SPM12 was applied to these gray matter probability maps, while covarying for age and total GMV (for more detail, see **Supplemental Text 2**). The resulting t-maps of XXY vs. XY differences in GMV were corrected for whole-brain comparisons using the revised version of AFNI’s 3dClustSim [41] to ensure family-wise error (FWE) < 0.05.

We assessed overlap between functional and structural findings by correlating the group difference t-score (XXY vs. XY) of seed-ROI rsFC in each ROI with the group difference t-score (XXY vs. XY) in regional GMV within that ROI. We also tested if the magnitude of inter-regional rsFC disruption in XXY was coupled with the magnitude of inter-regional GMV covariance disruption (see **Supplementary Text 3**) and found no statistical evidence of such a relationship.

### Inter-relating Variation in rsFC and Variation in Clinical Outcomes in XXY

To assess the role of the left DLPFC connectivity in explaining cognitive and behavioral variation in XXY, we separately conducted Pearson correlations of inter-individual variation in IQ and total CBCL score with inter-individual variation in each gray matter voxel’s connectivity with the left DLPFC in XXY. To identify regions showing significant correlations, we applied a cluster-size threshold to ensure full FWE < 0.05 by correcting each individual map to *p* <0 .025, accounting for our analysis of both CBCL and IQ (p < 0.002, cluster size threshold = 26 voxels). This thresholding was determined using the revised version of AFNI’s 3dClustSim [41].

## RESULTS

### Participant characteristics

Our study cohort included 75 XXY and 84 XY youth aged 6-25 years old (mean age 17 years). Groups did not significantly differ in terms of age (*p* = 0.45) or average head motion (*p* = 0.12). Mean IQ within the XXY group was 94.9 (SD = 12.13) and mean total CBCL score was 57.09 (SD = 9.20). For further details see **Supplementary Table 1**.

### XXY syndrome is associated with a distributed disruption of prefrontal connectivity

One 32-voxel cluster of increased global rsFC in the left DLPFC in the XXY compared to the XY group survived statistical thresholding and leave-one-out conjunction analyses (see **Methods, Figure 1**).

**Figure 1.**
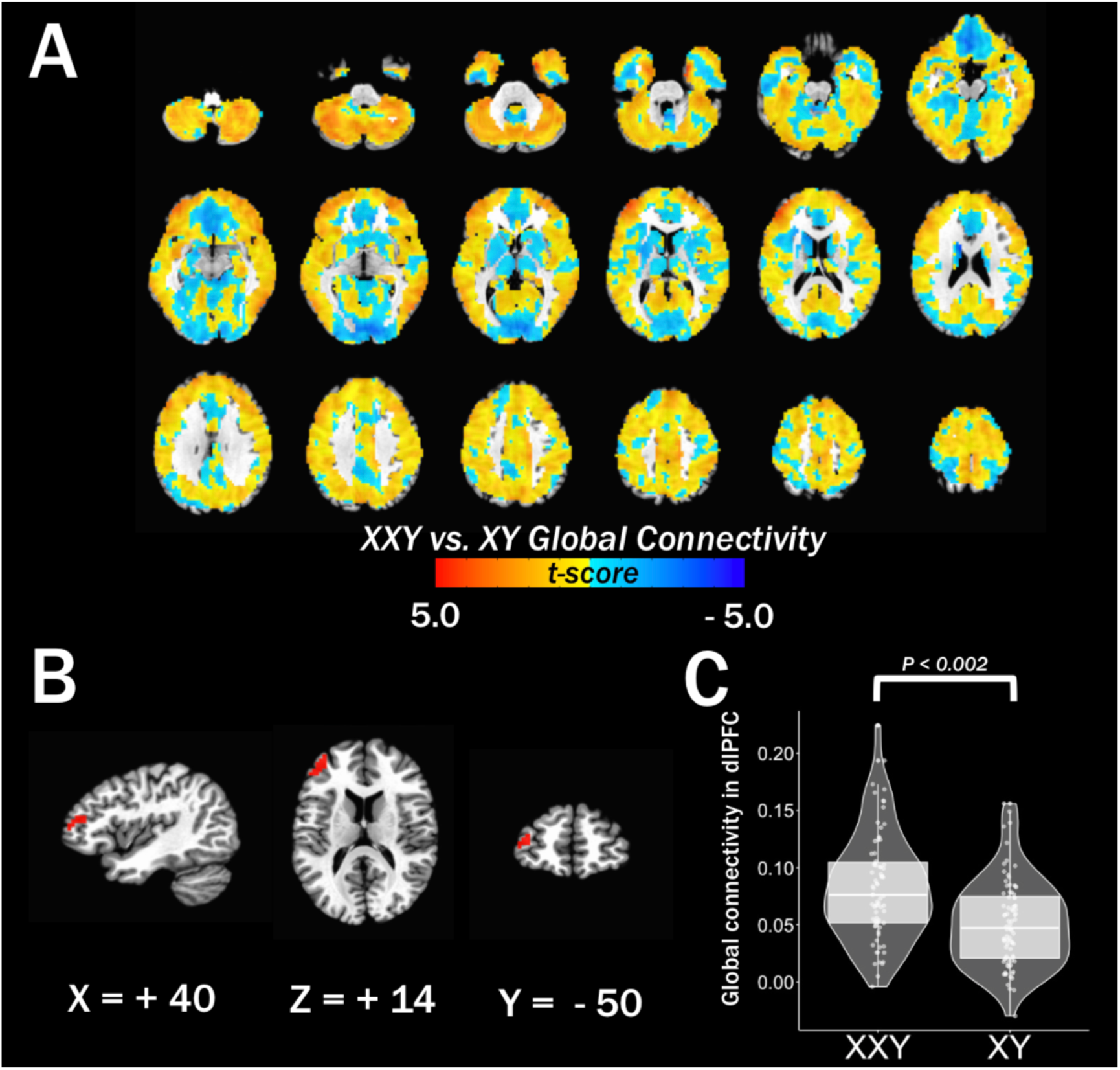
Increased dorsolateral prefrontal cortex global rsFC in XXY syndrome. **(A)** Unthresholded map of group differences (XXY-XY) in global connectivity while covarying for age and motion effects. (**B)** Significant 32 voxel cluster of increased global connectivity in the left DLPFC in XXY. Voxels survived all leave-one-out validation permutations at p = 0.002, cluster size threshold = 21 voxels). **(C)** Average global connectivity in left DLPFC voxels by group.

Seed-based analysis of group differences in voxel-wise connectivity with the left DLPFC revealed 24 clusters showing significantly increased left DLPFC connectivity in XXY syndrome compared to XY controls. Division of these distributed clusters by major anatomical divisions and labelling against an existing brain-wide rsFC parcellation (see **Methods**) yielded 39 regions of interest (ROIs) for downstream analyses (**Figure 2A, Table 1, Supplementary Movie 1**).

**Figure 2.**
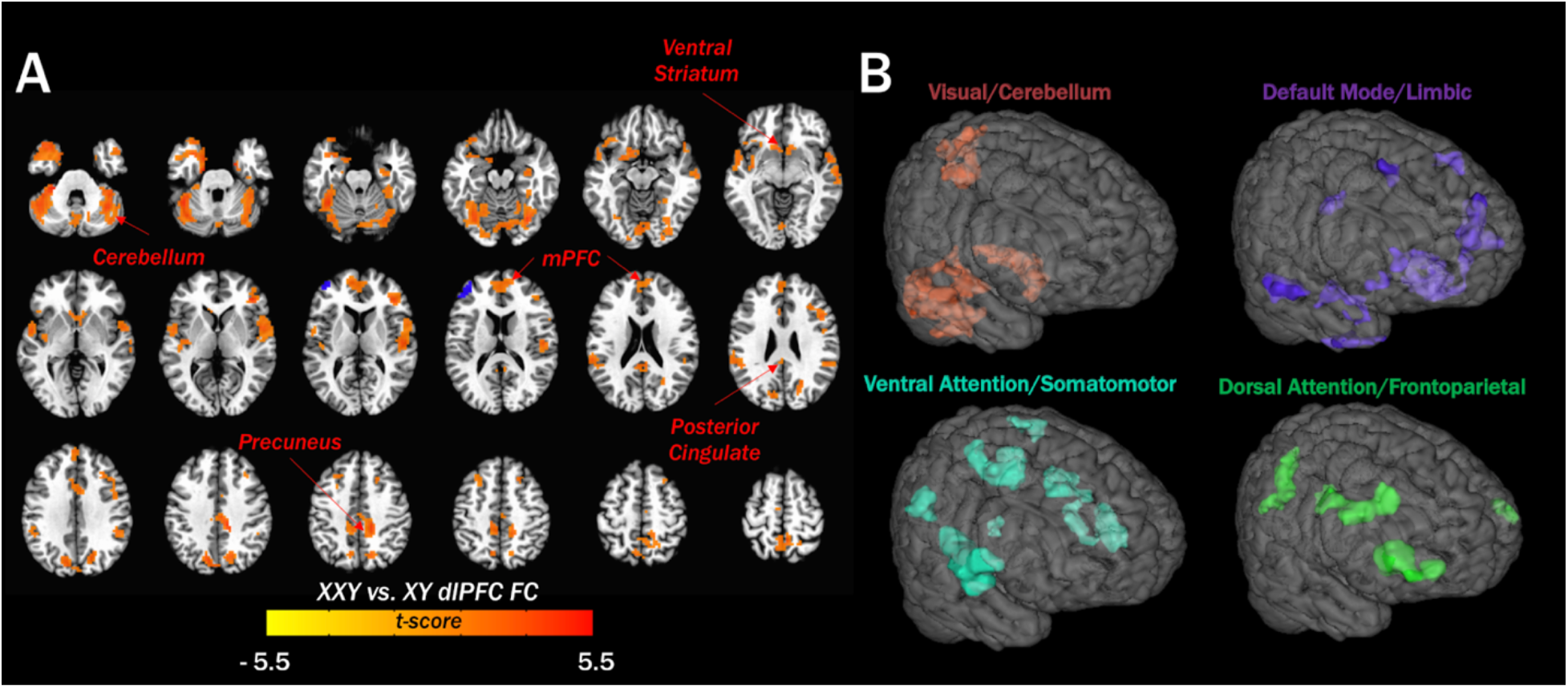
XXY syndrome increased rsFC between the left DLPFC and a distributed set of cortical and subcortical regions. **(A)** Regions showing significantly increased connectivity (in red) with the left DLPFC seed (in blue) in XXY syndrome relative to XY controls. **(B)** Volumetric visualization of regions showing increased DLPFC rsFC in XXY syndrome, grouped and colored according to their overlap with canonical resting-state FC networks in the Yeo-Krienen 7-network solution [45]: Red = Visual/Cerebellum, Purple = Default Mode/Limbic, Turquoise = Ventral Attention/Somatomotor, Lime Green = Dorsal Attention/Frontoparietal).

**Table 1.** ROIs showing increased dlPFC connectivity in XXY.

| ROI | Talairach Coordinates of peak signal |  |  | Voxels (3mm<sup>3</sup>) |
| --- | --- | --- | --- | --- |
|  | X | Y | Z |  |
| Left Cerebellum (VI) | 37 | 46 | -21 | 233 |
| Right Cerebellum (Crus) | -37 | 52 | -30 | 174 |
| Left Superior Medial Gyrus | 7 | -52 | 11 | 151 |
| Right Cerebellum (VIII) | -28 | 43 | -45 | 150 |
| Left Medial Temporal Pole | 31 | -13 | -33 | 135 |
| Left Cerebellum (X) | 25 | 31 | -33 | 125 |
| Left Superior Temporal Gyrus | 49 | 1 | 2 | 113 |
| Right Heschls Gyrus | -49 | 16 | 11 | 109 |
| Right Cuneus | -16 | 67 | 32 | 108 |
| Right Middle Cingulate Cortex | -13 | 34 | 38 | 99 |
| Right Cerebellum (VI) | -31 | 64 | -18 | 96 |
| Right Inferior Frontal Gyrus | -40 | -34 | 8 | 88 |
| Right Insula Lobe | -49 | -1 | 5 | 86 |
| Left Cuneus | 10 | 73 | 32 | 83 |
| Right Precuneus | -4 | 46 | 56 | 81 |
| Left SupraMarginal Gyrus | 55 | 34 | 23 | 76 |
| Right Inferior Frontal Gyrus | -43 | -13 | 26 | 76 |
| Right Olfactory Cortex/Caudate | -7 | -10 | -6 | 76 |
| Right Middle Temporal Gyrus | -61 | 13 | -6 | 63 |
| Right Inferior Temporal Gyrus | -37 | -1 | -36 | 61 |
| Right ParaHippocampal Gyrus | -19 | 4 | -24 | 61 |
| Left ParaHippocampal Gyrus | 16 | -1 | -24 | 58 |
| Left Heschls Gyrus | 40 | 19 | 8 | 55 |
| Left Posterior Cingulate Cortex | 4 | 37 | 23 | 52 |
| Left Cerebellum (VIII) | 4 | 61 | -33 | 44 |
| Right SupraMarginal Gyrus | -46 | 37 | 29 | 43 |
| Left Middle Cingulate Cortex | 4 | 37 | 44 | 42 |
| Left Inferior Frontal Gyrus | 31 | -25 | -15 | 42 |
| Left Supplementary Motor Area | 7 | 13 | 62 | 38 |
| Left Middle Frontal Gyrus | 22 | -25 | 41 | 37 |
| Right Anterior Cingulate Cortex | -4 | -7 | 29 | 35 |
| Left Middle Frontal Gyrus | 40 | -49 | 11 | 32 |
| Right Middle Frontal Gyrus | -22 | -19 | 44 | 29 |
| Left Cerebellum (VI) | 19 | 58 | -15 | 29 |
| Left Middle Occipital Gyrus | 22 | 79 | 8 | 23 |
| Right Superior Occipital Gyrus | -25 | 70 | 17 | 23 |
| Right Cerebellum (VIII) | -31 | 49 | -45 | 23 |
| Right Insula Lobe/Olfactory/Putamen | -28 | -10 | -6 | 22 |
| Left Lingual Gyrus | 10 | 55 | 0 | 21 |

*Note: ROI names determined by anatomical location of the voxel of greatest group difference in dlPFC connectivity according to the Eickhoff-Zilles macro labels in Talairach space [46].*

Clustering and annotating these ROIs against classical rsFC networks [45], (see **Supplementary Text 4**), revealed that altered global rsFC of the left DLPFC in XXY syndrome is driven by dysconnectivity between the left DLPFC and 4 distributed brain systems: [1. Visual/Cerebellum, 2. Default Mode/Limbic, 3. Ventral Attention/Somatomotor, 4. Dorsal Attention/Frontoparietal] (**Figure 2B, Supplementary Figure 1**). The left DLPFC seed fell within the Frontoparietal network according to this annotation. Permutation testing (see **Methods**) indicated that dysconnectivity amongst these ROIs was predominantly restricted to edges linked to the left DLPFC cluster of altered global rsFC in XXY syndrome, and some intra-cerebellar and parieto-cerebellar edges (**Supplementary Figure 2**).

### Changes in rsFC and regional anatomy are partially coupled in XXY syndrome

Voxel-wise analysis of group differences in GMV (controlling for age and total GMV) revealed relative volumetric expansion in parieto-occipital regions alongside relative volumetric reduction in cerebellum, subcortex, prefrontal cortex, and medial temporal lobes (*p* < 0.001, cluster size threshold = 306 1mm isotropic voxels; **Figure 3A, Supplementary Table 2**). To test the interrelationship between altered rsFC and altered GMV in XXY syndrome, we asked if inter-ROI variation in the magnitude of altered DLPFC connectivity was correlated with inter-ROI variation in the (unsigned) magnitude of GMV group differences (**Methods**). The magnitude of group differences in ROI rsFC dysconnectivity with the left DLPFC in XXY syndrome was positively associated with the magnitude of group differences in VBM using robust correlation (*r* = 0.47**; Figure 3B**). This correlation was statistically significant compared to a null distribution of correlations from group permutation (*p*_*perm*_ = 0.02; see **Supplementary Text 5**). Thus, brain regions showing greater dysconnectivity with the DLPFC in XXY syndrome also tend to be those that show greater group-level alterations of relative GMV.

**Figure 3.**
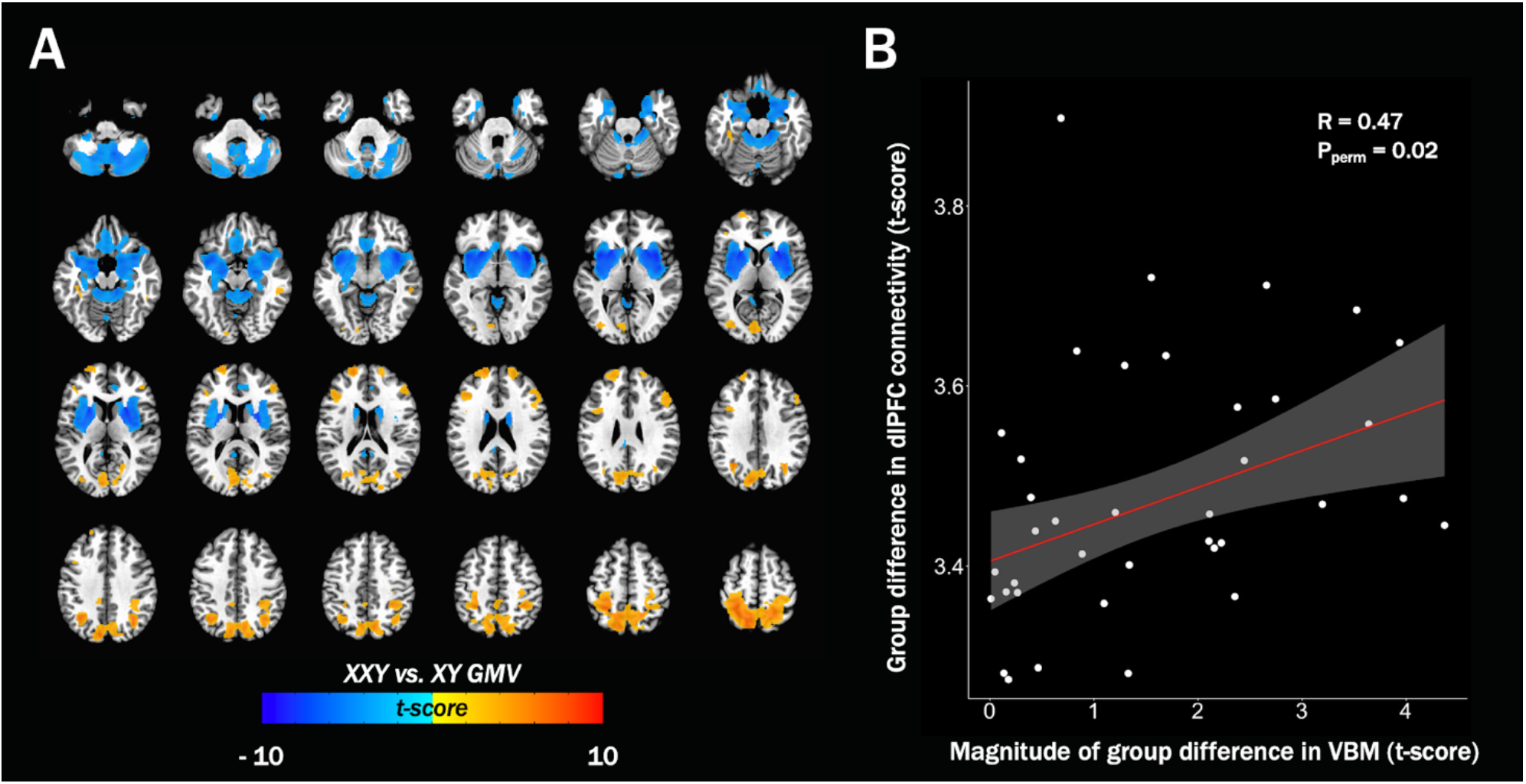
Altered regional brain volume in XXY syndrome and its relationship with rsFC changes. **(A)** Volumetric group differences in (XXY-XY; p < 0.002, cluster size threshold = 306 1mm isotropic voxels). **(B)** Scatterplot showing correlation between group difference in DLPFC connectivity (XXY-XY) and magnitude of group difference in VBM (XXY-XY) at each functionally defined ROI.

### Prefrontal connectivity is negatively correlated with psychopathology in XXY

Whole-brain voxel-wise correlations between inter-individual variation of left DLPFC rsFC and inter-individual variation in total CBCL score identified clusters of negative correlation in the left precuneus and in the bilateral inferior temporal gyri (ITG) (*p* < .002, FWE = 0.025; Peak *r* values: precuneus *r* = −0.45, Left ITG *r* = −0.48, Right ITG *r* = −0.43; **Figure 4A**). Conjunction with the map of altered DLPFC connectivity in XXY syndrome (**Figure 2A**) identified an overlap in the right precuneus -- indicating that this region not only shows increased connectivity with the DLPFC in XXY syndrome as compared to XY controls, but that also amongst individuals with XXY syndrome, greater DLPFC-precuneus connectivity is associated with less severe psychopathology. No clusters showed significant correlation between DLPFC connectivity and IQ. To better qualify our findings for rsFC-CBCL relationships in XYY syndrome, we also conducted the same analysis in the pediatric subset of XY controls with available CBCL data, and found that clusters identified in XXY syndrome were nonsignificantly positively correlated with total CBCL scores in euploidic male controls (see **Supplementary Text 6)**. Secondly, to gauge relative contributions of each CBCL subscale, we conducted sparse canonical correlation analysis (sCCA) between significant voxels in the XXY group and subscales of the CBCL. We did not observe that the correlation with total psychopathology scores was driven by specific subscales of the CBCL (see **Supplementary Text 6**).

**Figure 4.**
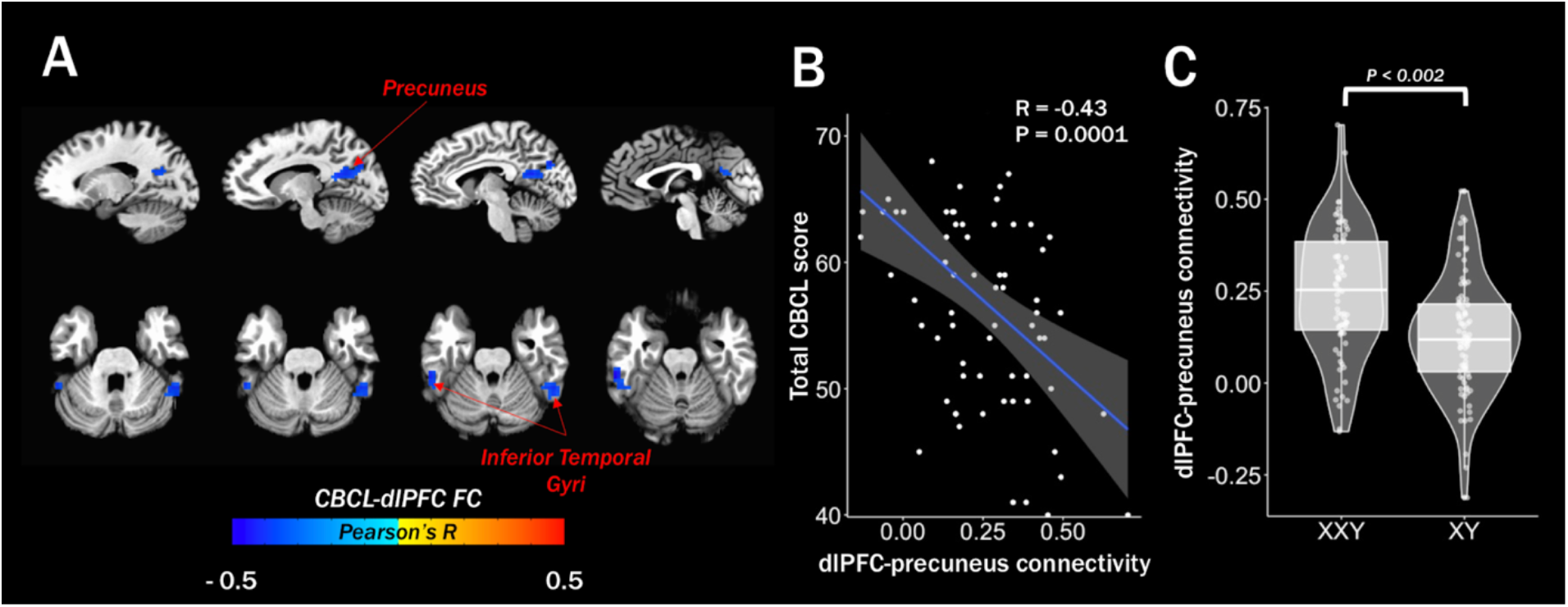
Relationships between DLPFC FC and psychopathology in XXY syndrome. **(A)** Regions showing significant correlations between DLPFC connectivity and total CBCL score in XXY syndrome (p < 0.002, cluster size threshold = 26 voxels, FWE = 0.025). **(B)** Scatterplot showing inverse correlation between DLPFC connectivity and total CBCL score among precuneus voxels that also show significant DLPFC increase in XXY. **(C)**. Boxplot showing average DLPFC connectivity in these precuneus voxels by group.

## DISCUSSION

Our study provides the first whole-brain screen for rsFC alterations in XXY, and their links with brain structure and behavioral measurements. We provide evidence for widespread disruption in functional brain organization in XXY that is partly coupled with anatomical alterations and related to the severity of overall psychopathology. These results inform our understanding of XXY syndrome as a medical condition in its own right, and as a genetics-first model for mechanisms of genetic risk in psychiatry more generally.

### Prefrontal dysconnectivity in XXY syndrome

We found significantly increased global connectivity in the left DLPFC in XXY which was associated with overconnectivity with a distributed set of cortical, subcortical, and cerebellar regions. These regions collectively encompass several major canonical rsFC networks within the human brain [45]. This pattern of altered rsFC in XXY syndrome connects with existing literature in several notable ways.

First, many of the regions showing overconnectivity with the DLPFC in XXY syndrome have been reported to show prefrontal underconnectivity in Turner syndrome (45, X) [DLPFC-intraparietal sulcus [11], DLPFC-angular gyrus [47]]. These inverted disruptions of DLPFC rsFC in XXY and Turner syndrome strengthen the case for a direct effect of X-chromosome dosage on frontoparietal rsFC organization of the human brain. Second, the nature of rsFC alterations observed in XXY syndrome within the current study shows a mixed pattern of partial overlap and divergence with those seen in other genetically defined neuropsychiatric disorders (specific comparisons provided in **Supplementary Text 7**). Consequently, an important goal for future work will be explicitly testing this hypothesis in prospectively designed comparative studies of rsFC across multiple neurogenetic disorders. Third, abnormalities of DLPFC rsFC, such as those we report here in XXY syndrome, have also been associated with diverse idiopathic neuropsychiatric disorders [48–51]. This observation builds on the occurrence of DLPFC rsFC alterations across diverse genetically defined neuropsychiatric disorders (see above) and may reflect the role of DLPFC in those higher-order domains of brain function [52] which are core to the manifestation of diverse psychiatric presentations. Given the position of the DLPFC towards the peak of the functional hierarchy [53], disruption may alter connectivity amongst the functionally disparate regions we observed. Finally, longitudinal studies of rsFC in typically developing youth have revealed substantial reorganization of DLPFC connectivity with other brain regions en route to adulthood [54, 55]. In this context, our findings in XXY syndrome could potentially be conceptualized as a shift in the timing of a normative developmental progression in rsFC, although a formal test of this hypothesis must await the accumulation of later longitudinal rs-fMRI datasets in XXY syndrome.

### Partial convergence of rsFC and anatomical changes in XXY syndrome

For regions showing altered rsFC with the DLPFC in XXY syndrome, we observed that interregional variation in the magnitude of rsFC dysconnectivity was correlated with the absolute magnitude of regional brain volume change -- indicating that disruptions of rsFC in XXY syndrome may be mechanistically linked to disruptions of anatomy that are visible *i*n vivo. Our study design (and observational methods in humans generally) prevents direct analysis of causality, although longitudinal data could potentially support a causal relationship. It is also possible that XXY syndrome induces a coordinated disruption of both these brain phenotypes without one necessarily driving the other. Interestingly, in vivo imaging has also detected overlapping alterations of brain structure and function in other genetically defined [e.g. 22q11 copy number variations, [19]] and idiopathic [e.g. ASD [56] and attention-deficit/hyperactivity disorder [57]] neuropsychiatric disorders. Thus, although such overlaps are certainly not a general rule across psychiatric disorders [58, 59] -- our findings motivate broader multimodal analyses of regional brain disruption on neuropsychiatric disorders.

### Severity of psychopathology in XXY correlates with variation in DLPFC rsFC

We observed that variation in total psychopathology, but not in IQ, correlates with DLPFC rsFC in XXY syndrome. Strikingly, comparison with prior work suggests that prefrontal dysconnectivity is a correlate of psychopathology among multiple neurogenetic conditions, idiopathic psychiatric disorders, and in healthy populations [14, 16, 20, 21, 60, 61]. We find DLPFC rsFC correlates with total psychopathology in XXY syndrome within the inferior temporal gyrus (ITG) and the precuneus. The functioning of the ITG has been associated with higher order social communication [62] and visual processing [63]. The precuneus is a key node within the brain’s default mode network and implicated in several higher order capacities including working memory and self-directed cognition [64]. The precuneus focus where DLPFC rsFC connectivity correlates negatively with psychopathology in XXY is notable for also showing increased DLPFC rsFC in XXY syndrome as compared to XY controls. Thus, the “direction” of group differences and within-group symptom correlation for DLPFC-precuneus rsFC in XXY syndrome appear to be opposing. This preliminary finding awaits replication in independent cohorts but could potentially represent a compensatory process such as has been reported in other disorders [65, 66]. Importantly, the pattern of CBCL correlation with this edge in XYY syndrome was both qualitatively and quantitatively distinct from that in the XY group and contributed to by multiple CBCL subscales. An important caveat is that CBCL data was only available on XY participants under 18 (N=45), therefore we cannot rule out confounding age differences in this analysis.

### Limitations and future directions

Our findings should be considered with a set of important caveats and limitations in addition to those already mentioned. Firstly, as our sample is not population based, we cannot assume that the phenotype of XXY in this sample is fully representative of all cases of XXY. Indeed, as a large number of individuals with XXY go undiagnosed, it is possible that those with higher penetrance are more frequently diagnosed, and therefore a more extreme phenotype is overrepresented in our sample. This is a known problem in the SCA literature and clinical studies of genetically defined neuropsychiatric disorders more broadly [5, 67, 68]. Second, carriage of a supernumerary X-chromosome in males is also associated with hypogonadism and testosterone deficiency beginning at the onset of puberty [69]. Future research should assess the degree to which hormonal influence could act as a mediating variable for SCA on brain function. Third, recent studies have argued that substantially larger sample sizes are necessary to reliably correlate brain function with behavior [60, 70]. Collecting data from such a large sample is infeasible when studying rare neurogenetic conditions such as XXY. Therefore, future research should seek to independently reproduce brain-behavior correlations observed here to ensure their reliability.

### Summary

Notwithstanding these limitations, our study provides the first systematic investigation of brain connectivity in XXY syndrome -- which represents a common and clinically significant genetic risk for neuropsychiatric impairments. In doing so, we identify a distributed pattern of altered prefrontal rsFC in XXY syndrome compared to XY males. These changes in brain connectivity are partly correlated with regional anatomical disruptions in XXY syndrome, and also overlap with regions where variation in prefrontal connectivity is correlated with variation in psychopathology amongst affected individuals. Taken together, these findings provide a fuller understanding of functional and structural brain changes in XXY syndrome as a medical disorder in its own right, and as a “genetics-first” model for neuropsychiatric impairment more broadly.

## ACKNOWLEDGEMENTS

The authors would like to thank the patients and their families for their participation in this study and the Association for X and Y Chromosome Variations (AXYS; https://genetic.org) for their assistance in recruitment efforts. All research was funded by the NIMH Intramural Research Program. (Clinical trial reg. No. NCT00001246; clinicaltrials.gov; NIH Annual Report Number, ZIAMH002949-03; Protocol number: 89-M-0006).

## CONFLICT OF INTEREST

The authors declare no conflicts of interest.

## Notes

### Competing Interest Statement

The authors have declared no competing interest.

